# The chicken chorioallantoic membrane model for isolation of CRISPR/cas9-based HSV-1 mutant expressing tumor suppressor p53

**DOI:** 10.1101/2023.05.12.540548

**Authors:** Mishar Kelishadi, Hosein Shahsavarani, Alijan Tabarraei, Mohammad Ali Shokrgozar, Ladan Teimoori-Toolabi, Kayhan Azadmanesh

**Affiliations:** Department of Molecular Virology, Pasture Institute of Iran, Tehran, Iran; Department of Cell and Molecular Biology, Faculty of Life Science and Biotechnology, Shahid Beheshti University, Tehran, Iran; Laboratory of Regenerative Medicine and Biomedical Innovations, Pasteur Institute of Iran, National Cell Bank, Tehran, Iran; Infectious Diseases Research Center, Golestan University of Medical Sciences, Gorgan, Iran; Department of Virology, Faculty of Medicine, Golestan University of Medical Sciences, Gorgan, Iran; Molecular Medicine Department, Biotechnology Research Center, Pasteur Institute of Iran, Tehran, Iran

## Abstract

Oncolytic viruses (OVs) have emerged as a novel cancer treatment modality, which selectively target and kill cancer cells while sparing normal ones. Among them, engineered Herpes Simplex Virus type 1 has been proposed to be employed as a potential treatment of cancer and was moved to phase III clinical trials. In this study, to improve oncoselectivity and oncotoxicity properties, the UL39 gene of the ICP34.5 deleted HSV-1 was manipulated with the insertion of the EGFP-p53 expression cassette utilizing CRISPR/ Cas9-mediated editing genome. The ΔUL39/Δγ34.5/HSV1-p53 mutant was isolated using the chorioallantoic membrane (CAM) of fertilized chicken eggs as a complementing membrane to support the growth of the viruses with gene deficiencies. Phenotypic characterization of ΔUL39/Δγ34.5/HSV1-p53-infected cells was compared with the parent Δγ34.5/HSV-1 in vitro. Our results indicate that the CAM model can be a promising strategy for isolating recombinant virus such as HSV-1-P53 that is unable to replicate in cell lines due to the death induced by exogenous p53 during virus replication.

## Introduction

Today, cancer is a global health problem responsible for one death in six worldwide [1]. Poor overall survival of patients affected with advanced types of cancer and resistant to conventional therapies such as surgery, chemotherapy, and radiotherapy, indicate a great need for the development of novel therapeutic strategies [2].

HSV-1-based recombinant viruses emerged as a new platform in the development of oncolytic viruses due to their capacity for foreign genes, efficient replication, broad host cell range, and safety because their genome is not incorporated into the human genome [3–6].

Human alphaherpesvirus 1 (HSV-1) is a ubiquitous eukaryotic pathogen, belonging to the Herpesviridae family and to the Alphaherpesvirinae subfamily, that has a relatively large enveloped virus with a 152-kb linear double-stranded genome and codes about 80 proteins, half of which are not essential for virus replication [6–8]. The UL39 (ICP6) gene encodes the large subunit of HSV-1 ribonucleotide reductase, a protein complex that converts ribonucleotides to deoxyribonucleotides and provides a major pathway in the synthesis of DNA precursors for its replication. However, it is not crucial for viral growth in dividing cells but its function is required for viral replication and DNA synthesis in quiescent or serum-starved cells as well as neuronal cells. This enzyme is overexpressed in dividing cells such as tumor cells; as a consequence, an ICP6*-*null mutant preferentially replicates in human tumor cells but not in normal cells with limited dividing activity [9–11].

The superior characteristic of HSV-1, especially oncolytic herpes simplex virus-1 which lacks ICP34.5 is its ability to induce cell death via apoptosis-dependent and independent mechanisms in a cell-specific manner in which caspase-8 is crucial for HSV-1-induced apoptosis [12–17].

Studies of HSV-dependent apoptosis have resulted in conflicting data. However, the recent consensus that viral factors such as the US3, US5, ICP4, ICP6, ICP22, ICP27 proteins, glycoprotein D, glycoprotein J, and the latency-associated transcript (LAT) are responsible for the anti-apoptotic activity, ICP0 and ICP27 are multifunctional proteins which regulate many aspects of the cellular and viral functions including apoptotic responses [10-12, 14-16, 18].

Although ICP0 in most studies, has been identified as a pro-apoptotic HSV-1 protein, a few studies have reported that ICP0 can act as an E3 ubiquitin ligase to induce efficient degradation of the p53 protein and inhibit the infected cells from p53-mediated apoptotic responses [10, 16–19].

Together, given the importance of apoptosis in the HSV-1 life cycle, careful timing of apoptosis induction by the viral genes of HSV-1 is crucial in designing therapeutic modality from this virus [16, 17, 20, 21].

Previous studies have supported the pivotal role of p53 in apoptosis during HSV1-induced oncolysis. P53 is a regulatory protein and a nuclear transcription factor that plays a key role in controlling cell division and cell death. In unstressed cells, the expression of p53 is normally maintained at a low level through ubiquitination and proteasome-mediated degradation of this protein [18, 22]. A variety of cellular stresses including DNA damage, hypoxia, oncogene activation, and viral infections lead to the stabilization of p53. In this regard, through increased half-life of p53 and transcriptional activation of p53-responsive genes DNA repair or apoptosis are enhanced and the propagation of cells with serious DNA damage is inhibited [18, 22, 23].

An extensive search revealed that p53 mutations are present in approximately 50% of cancers [20, 22, 24, 25]. As well, many cancer cells have a nonfunctional p53 pathway which results in defects in the apoptosis pathway. Anyhow, the functional status of p53 in cancer cells may alter the mechanism of cell death and affect the chemo-resistant phenotype of these cells [20, 22, 24, 25].

Based on reported studies, exogenous expression of p53 in human cancer cells during replication of oncolytic viruses such as stomatitis virus (VSV), Newcastle disease virus (NDV), and adenovirus enhance the cell death leading to anti-tumor effects of these viruses [25–28].

Although HSV-1 mutants were suggested to be a promising candidate for sensitizing the radiotherapy/chemotherapy-resistant tumors [12], some cancer cells (e.g. MCF7 cells) are resistant to HSV-1-dependent apoptosis [10]. Therefore, it was hypothesized that restoration of wild-type P53 activity by HSV-1 oncolytic would be a potential approach for triggering the p53-mediated pro-apoptotic and enhancing the oncolytic potency in advanced tumors.

The CRISPR/Cas9 (Clustered regularly interspaced short palindromic repeats/CRISPR-associated 9) system is a complex of a single guide RNA (sgRNA) that target particular genomic loci and a Cas9 as an endonuclease that makes a double-strand DNA break (DSB). This cleaved DNA subsequently can induce deletions and mutations at the target site or incorporate a transgene into these sites by homologous recombination [29].

In the effort to employ HSV-1 as a therapeutic modality and improve the oncoselectivity and oncotoxicity of this virus, we aimed to inactivate the UL39 gene of a mutant HSV-1 which lacks both copies of the γ34.5 gene which through the insertion of a fluorescent P53GFP fusion gene expression cassette using the CRISPR-Cas9 system. We attempted to isolate the recombinant virus from Vero and CAM. In the next step, we evaluate the oncolytic property of ΔUL39/Δγ34.5/HSV1-p53.

## Materials and Methods

### Cell Lines, Viruses, Plasmids and Chicken eggs

In this study, Vero (African green monkey kidney, NCBI-C101), BHK-21 (Baby hamster kidney, NCBI-C107), A549 (human lung epithelial, NCBI- C137), MDA-MB-468 (Human Adenocarcinoma, NCBI- C208), Hela (Human cervical carcinoma, NCBI- C115), HEK 293 (Human Embryo Kidney, NCBI- C497), HEK 293T (Human Embryonic Kidney, NCBI- C498), Caco-2 (Human Colorectal Adenocarcinoma, NCBI- C139), and NIH3T3 (Mouse Embryo cell, NCBI- C156) cell lines were obtained from the National Cell Bank of Pasteur Institute of Iran. These cells were cultured in high glucose Dulbecco’s modified Eagle’s medium (DMEM) (Gibco, Germany) supplemented with 10% heat-inactivated Fetal Bovine Serum (FBS; Gibco), 2mM L-glutamine, and 1 % Penicillin/Streptomycin (Gibco, Germany) at 37°C with a humidified atmosphere containing 5% CO2. All cell lines were tested to be free from the contaminations.

The principal virus used in this study was Δγ34.5/HSV-1 virus (Herpes simplex virus1 Red-Red), an ICP34.5-null mutant in which both copies of ICP34.5 were replaced by an insert carrying a Blecherry reporter gene driven by the cytomegalovirus promoter [30, 31].

pcDNA3.1 (+) IRES GFP plasmid (Addgene, #51406), sgRNA/Cas9 cloning vector pX459-puro (Addgene, *#*62988), sgRNA/Cas9 cloning vector pX459-mCherry (Addgene, *#*64324), the pIRES2-EGFP-p53 WT Plasmid (Addgene, #49242)

Ten-day-old Specific Pathogen Free (SPF) embryonated chicken eggs were purchased from Razi Vaccine & Serum Research Institute (Karaj, Iran). As it is generally accepted that the embryo cannot feel pain until approximately day 19, special permission for animal experiments was not required [32–37].

### Δγ34.5/HSV-1 preparation and DNA extraction

Vero cells were infected with Δγ34.5/HSV-1 at a multiplicity of infection (MOI) of 0.01. After 2 days, the cells were harvested after observation of the total cytopathic effect. The supernatant was titrated, aliquot, and stored at −80°C.

DNA was purified from virus-infected Vero cells stock by a commercially available kit (High Pure Extraction Kit; Roche Diagnostics GmbH, Mannheim, Germany) according to the manufacturer’s instructions.

### Design and cloning gRNA oligo’s into Cas9 vector

The CRISPR/Cas9 system used in this study was constructed by introducing synthesized oligo primers targeting the ul39 gene of Δγ34.5/HSV-1 into sgRNA/Cas9 cloning vector pX459 according to Feng Zhang’s lab recommendations: (https://www.addgene.org/crispr/zhang/). Briefly, the gRNAs were designed and selected using available online tools (http://crispr.mit.edu) and (http://www.rgenome.net/cas-offinder) to select the optimal sequence for maximizing double-stranded breaks (DSBs) while minimizing the off-target effect.

Two complimentary oligodeoxynucleotides gRNA F and gRNA R (Table 1) were annealed in a thermocycler and the resulting dsDNA fragment was then ligated into the *Bbs*I (ThermoFisher Scientifc, USA) site of linearized pX459. The resulting plasmid is named Cas9/gRNA_UL39._ The insertion of the gRNAs was confirmed with the test primers (CMV_P_F and gRNA R) (Table 1) and sequencing.

**Table 1.**
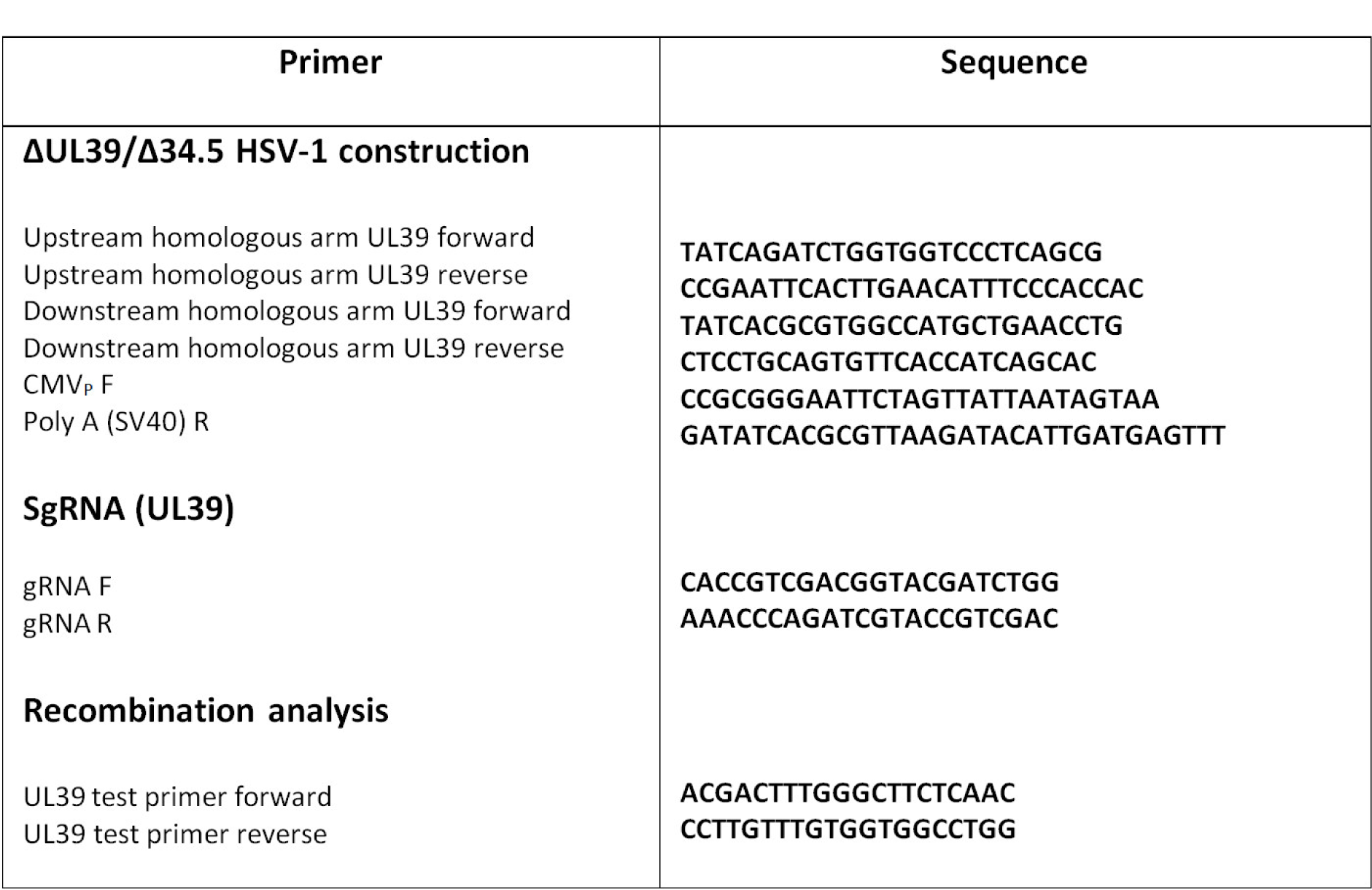
Primer sequences for generating homologous fragments of UL39 and recombination analysis.

### Construction of UL39 shuttle donor vector for homologous recombination (HR)

UL39 shuttle donor vector was constructed based on pJET1.2/blunt Cloning Vector backbone, a high-efficiency TA cloning vector (CloneJET PCR Cloning Kit, ThermoFisher Scientific, USA). Briefly, the full sequence of the expression cassette containing the CMV promoter, EGFP coding region, and the SV40 early mRNA polyadenylation signal was amplified from pcDNA3.1(+)IRES GFP using the forward primer (CMV_P_ F) containing *Eco*RI restriction site and the reverse primer (PolyA R) containing *Mlu*I (ThermoFisher Scientifc, USA) restriction site.

The resulting PCR product was purified using GeneJET PCR Purification Kit (Thermo Fisher Scientific, USA) according to the manufacturer’s manual and directly cloned in the pJET1.2/blunt cloning vector to generate GFP-pJET plasmid (Table1).

Homologous right and left fragments encoding about 856 and 823 nucleotides of the genome which flank the specific regions within the UL39 gene of Δγ34.5/HSV-1 genome were amplified by PCR with the primers; Upstream homologous arm UL39 forward) containing *Bgl*ll (ThermoFisher Scientifc, USA) restriction site and Upstream homologous arm UL39 reverse containing *Eco*RI (ThermoFisher Scientifc, USA) restriction site for Ul39 R fragment and with the primers Downstream homologous arm UL39 forward containing *Mlu*I restriction site and the reverse primer Upstream homologous arm UL39 reverse containing *Pst*I (ThermoFisher Scientifc, USA) restriction site for UL39L fragment. After purifying, the fragments were sub-cloned sequentially into the GFP-pJET at *Mlu*I/*Pst*I and *Bgl*ll/*Eco*RI sites. The resulting plasmid, which is also named pJET-UL39R-CMV-GFP-Ul39L was confirmed by Sanger sequencing (Table 1).

The p53 coding sequence in the pIRES2-EGFP-p53WT plasmid was cut out with *Nhe*I (ThermoFisher Scientifc, USA) and *Not*I (ThermoFisher Scientifc, USA) and ligated into the similar restriction sites of pJET-UL39R-CMV-GFP-Ul39L shuttle plasmid to generate pJET-UL39R-CMV-GFP-P53-Ul39L.

### Plasmids extraction and purification

Plasmid isolation and DNA fragments purification was performed using the ThermoScientific GeneJET Plasmid Miniprep Kit (Thermo Scientific, Waltham, MA USA) and GeneJET Gel Extraction Kit (Thermo Scientific, Waltham, MA USA), respectively according to the manufacturer’s instructions.

### Generation of ΔUL39/Δγ34.5/HSV1-p53 mutant using the CRISPRCas9 system

Transient transfections were carried out on BHK cells using ScreenFect™ A plus (Fujifilm WAKO, Japan) according to the manufacturer’s protocol. The reagent consisted of 1µg of DNA and 1µL of reagent per each well of a 24-well plate.

SgRNA/Cas9 cloning vector pX459-mCherry (Cat no. 64324; Addgene) and pJET-UL39R-GFP-p53-Ul39L plasmid were used as controls in all transfection experiments for monitoring the cell viability along with the reporter gene expression.

Briefly, BHK cells were seeded into 24-well plates (SPL Life Sciences, Korea) at a density of 0.05 x 10^6^ cells/well to obtain 80% confluency for transfection. Cells were then transfected with Cas9/gRNA_UL39_ plasmid that contains puromycin-resistance gene in the presence of 1 μM SCR7, a non-homologous end joining (NHEJ) inhibitor (Sigma-Aldrich, USA). Cas9 and gRNA were allowed to be expressed for 24h. The second transfection was done with pJET-UL39R-GFP-p53-Ul39L plasmid supplemented with 10 ug/mL of puromycin (Bio Basic, Canada). After 48h, inoculation with Δγ34.5/HSV-1 was done at the MOI of 1 in an incubator (37 °C with 5% CO2) for 1-2 h. The inoculums were then removed and the cells were gently washed with the pre-warmed PBS to remove any un-adsorbed input virus. The transfection/infection supernatant from BHK cells was harvested at 24–48 h, upon the appearance of the 80% cytopathic effect of the cell monolayer and after three freeze-thaw cycles, they were stored as aliquots at −80°C (Fig 1).

**Fig 1.**
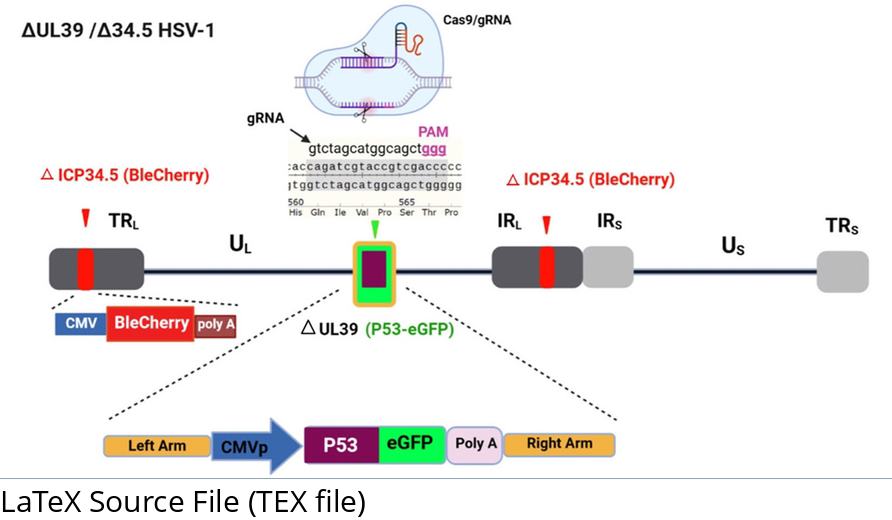
Schematic structure of ΔUL39(green)/Δ 34.5(red) HSV1-p53 that was developed in an ICP34.5 deleted HSV-1 backbone. Modifications include deletion of the UL39 gene and insertion of an EGFP-P53 expression cassette utilizing a CRISPR/Cas9-mediated editing genome system. The internal ribosome entry site (IRES) sequence between p53 and GFP genes allows the co-expression of the genes and facilitates their detection in mammalian cells while retaining the properties of wild-type p53.

### Isolation of the ΔUL39/Δγ34.5/HSV1-p53 mutant in vitro

96-well plated were seeded with 10^4^ Vero and BHK-21 cells per well, one day before infection. To isolate the recombinant viruses, confluent monolayers of cells were infected with different dilutions of the mutant viral supernatant. The plates were monitored daily using an inverted fluorescent microscope for EGFP expression at 1– 3 days post-infection.

### Isolation of the ΔUL39/Δγ34.5/HSV1-p53 mutant in CAM of the fertilized chicken eggs

For isolating the recombinant virus in the CAM, after checking the eggs for viability and creating a false air sac and a ‘window’ opening in the shell, the CAMs were inoculated with 0.1 ml each of serially 10-fold diluted viruses (The transfection/infection supernatants) [36, 38–40].

The opening was then sealed with povidone iodine-impregnated paraffin waxes to avoid contamination. The eggs were incubated in a Small Egg incubator Easy Bator 3 (Eskandari Industrial Group, Iran) at 34.5 °C and 58-60% humidity without rotation. After 72 h and before harvesting the CAMs, the eggs were kept in a refrigerator at 4^○^C for 2 h to constrict the blood vessels [41, 42]. The harvested membranes were placed in a petri dish containing normal saline supplemented with 4% penicillin/streptomycin to flatten the rolled CAM. Then the medium was gently removed. The pocks were analyzed using an inverted fluorescent microscope for eGFP or BleCherry expression.

For purification of the HSV1-P53 mutant, the pocks which were positive for both BleCherry and GFP signals (pocks containing the recombinant green/red virus) were isolated. Thereafter, they were dispersed by trypsin and after three freeze-thaw cycles, they were sub-cultured in the new CAM (Fig 2).

**Fig 2.**
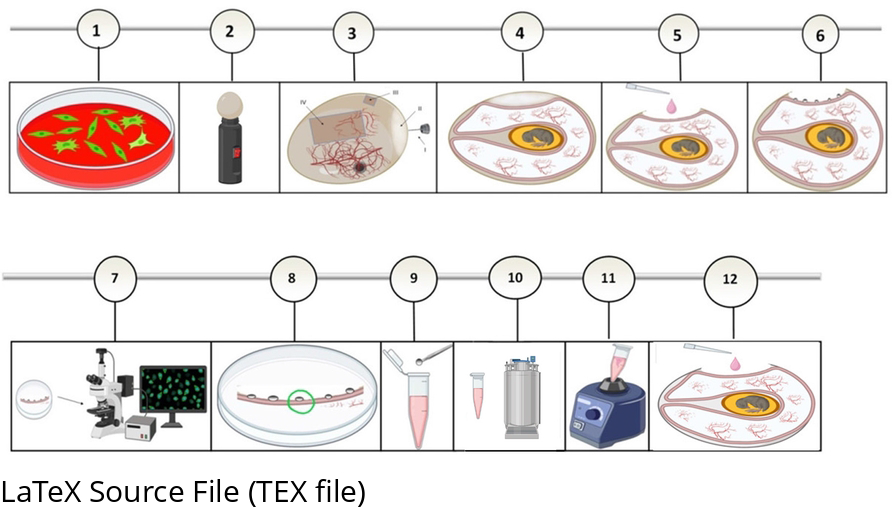
Experimenct workflow for construction of recombinant virus using chorioallantoic membrane (CAM) of fertilized chicken eggs. the process of transfection/infection of cells (1.2). Candling egg for visualizing the air sac (II) and identification of the egg vasculature (2.2). Preparation of artificial air sacs including the major steps; markings (I, III,IV), drilling (I) and creation of a square opening (III) and an operating window (IV) in the egg shell (3.2). Generation of the artificial air sac (4.2). Inoculation of the diluted virus onto the CAM membrane (5.2). The established pocks on the CAM membrane three days after the virus inoculation (6.2). Analyzing of the pocks using an invert fluorescent microscope for fluorescence protein expression (7.2). Selecting of the pock with fluorescence signals (8.2). Dipping the selected pocks into the culture medium (9.2). Three freeze*–*thaw cycles (10.2). Vigorous vortexing of the media containing the pock (11.2). Inoculation of the pock (virus) onto a new CAM (12.2).

### PCR analysis and sequencing for verification of homologous recombination

To verify recombination, the pocks positive for both BleCherry and GFP signals (pocks containing the recombinant green/red virus) were isolated for DNA extraction using High Pure Viral Nucleic Acid Kit (Roche Diagnostics GmbH, Mannheim, Germany) according to the manufacturer’s instructions. PCR was performed using a PCR master mix kit (Taq DNA Polymerase Master Mix RED 2x, Ampliqon, Denmark) in a total volume of 25 containing of 12.5 μl Taq DNA Polymerase, 1x Master Mix RED, (_∼_100–150 ng) of HSV-1 DNA and 0.2 μM of each forward and reverse test primers. The sequences of primers are given in Table 1. Program of PCR was as follows: 95 °C for 5 min; 30 cycles of 95 °C for 60 s, 55°C for 30 s, and 72 °C for 2 min followed by the final extension step at 72°C for 5 min. The PCR products were loaded on a 1% agarose gel and visualized by exposing it to ultraviolet (UV) light.

### Phenotypic Characterization of ΔUL39/Δγ34.5/HSV-p53mutant

Vero, BHK-21, A549, MDA-MB-468, Hela, HEK 293, HEK 293T, Caco-2, and NIH3T3 cell lines (10^4^ cell/well) were seeded into 96-well plates to characterize the newly isolated oncolytic virus phenotypically. They were infected the following day with 0.1 ml of various dilutions of the recombinant virus. After 1 hour of incubation at 37°C, cells were cultured in DMEM medium (containing 1% FBS) and were incubated for up to 5 days for GFP expression. All assays were performed in triplicates.

## Result

### Construction of ΔUL39/Δγ34.5/HSV-p53 and verification of homologous recombination

For the construction of the recombinant virus, firstly the sgRNA targeting the specific regions within the UL39 gene of the Δγ34.5/HSV1 was cloned into sgRNA/Cas9 cloning vector pX459.

The donor vector containing the UL39R-GFP-p53-Ul39L fragment was used for homologous recombination to improve the efficiency of homology-directed repair (HDR). Both Cas9/gRNA_UL39_ and the donor plasmids were transfected into BHK-21 cells.

The viral supernatant containing HSV-1 mutant was inoculated in Vero and BHK-21, MDA-MB-468, Hela and HEK 293T cell lines to isolate the recombinant virus but in all of them, the recombinant virus replication was limited to a single cell (Fig 3). So the viral supernatant was inoculated onto CAM (Fig 4).

**Fig 3.**
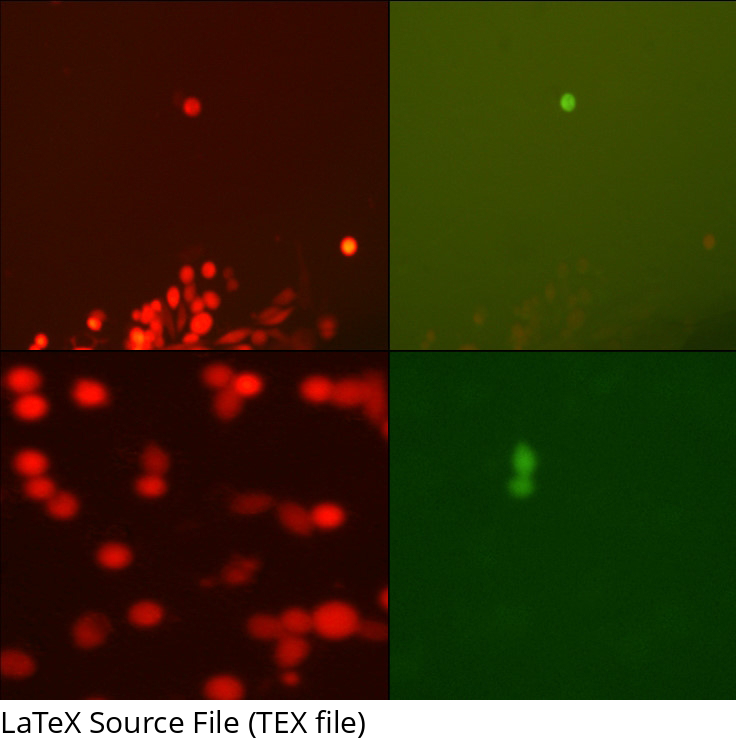
Representative results of inoculation of the viral supernatant that were expected to contain mutated viruses in BHK-21 (a, b) and MDA-MB-468 (c, d) cell lines. the recombinant virus replication was limited to a single cell (Magnification × 200).

**Fig 4.**
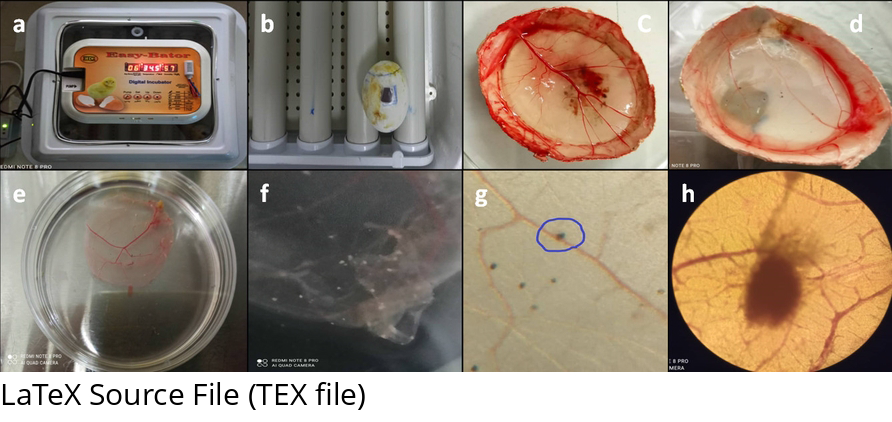
Representative results of using chorioallantoic membrane (CAM) of fertilized chicken eggs for construction of the recombinant virus. Incubation of the eggs in small egg incubator Easy Bator (a). The generated operating window in the egg shell (b). The harvested CAMs three days after inoculation of the virus with a titer of 10^4^ pfu/ml (c) and 10^2^ pfu/ml (d), illustrating visible pocks. The isolated membrane from the egg shell in a petridish (e). Macroscopic picture of pocks formed on CAM (f). Microscopic picture of pocks formed on CAM (g)(Scale bar: 40 µm). Higher magnification of the pock (h)(Scale bar: 200 µm).

Of note, high titer of the virus in the inoculum (10^7^ pfu/ml) induced no discrete pocks in CAM and the CAM membrane looked normal but fluorescent analysis of red Δ34.5/HSV-1 illustrated red confluent lesions in the CAM, reflecting the virus distribution throughout the CAM (Fig 5).

**Fig 5.**
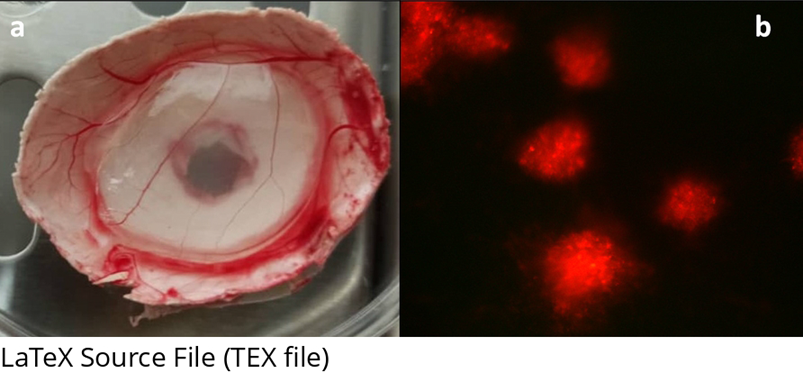
High titer of the virus resulted in confluent lesions and induced no discrete pocks in CAM. Macroscopic picture of high titer virus-infected CAM (a). Microscopic picture of high titer virus-infected CAM (b) (Magnification × 100).

Three days later, the pocks positive for both BleCherry and GFP signals were confirmed by visualizing by the inverted fluorescent microscopy (Fig 6) and molecular analysis (Fig 7).

**Fig 6.**
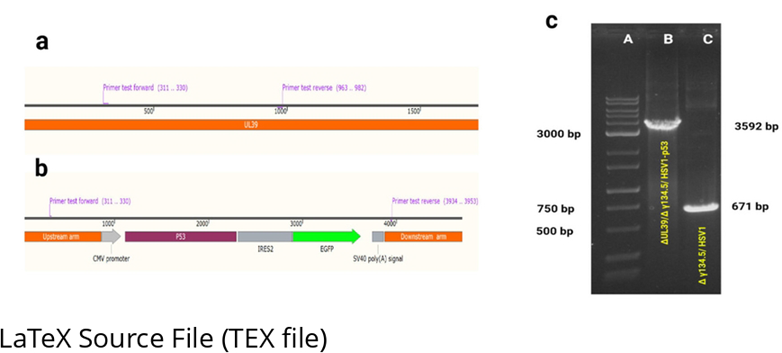
Fluorescence and brightfield images of representative pocks showing the EGFP & BleCherry signals confirming construction of the ΔUL39 (green)/Δ 34.5 (red) HSV1-p53 in chorioallantoic membrane (CAM) of fertilized chicken eggs (Magnification × 200).

**Fig 7.**
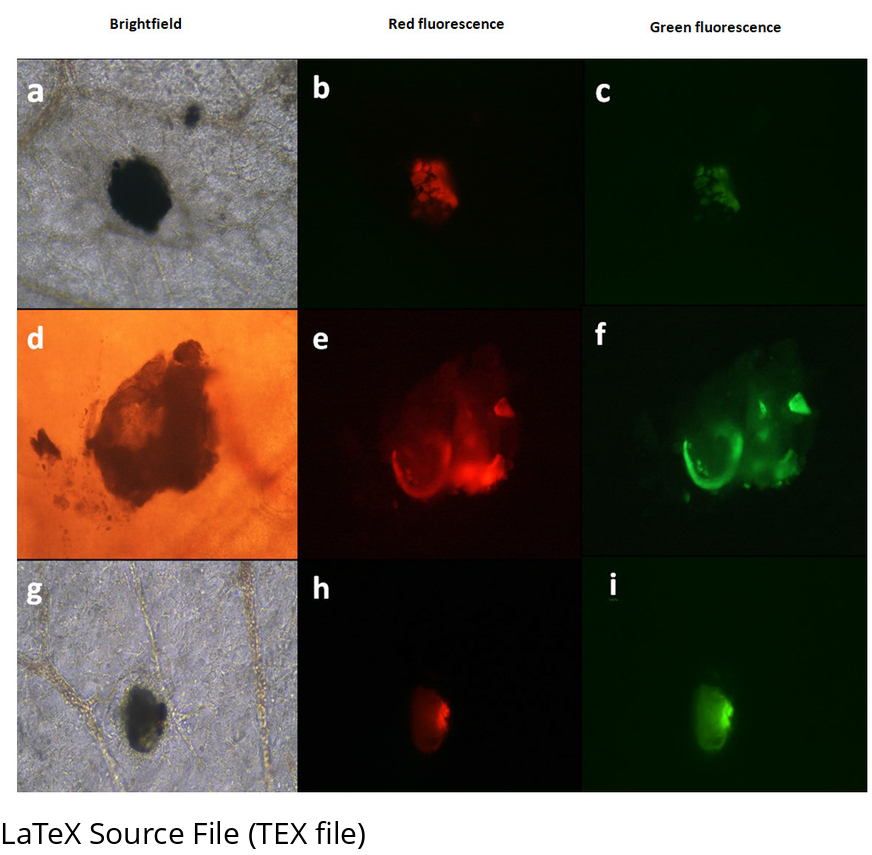
Schematic illustration of the test primers binding site to UL39 in. **Δ34.5**/HSV-1 parental virus (a) recombinant ΔUL39/Δ34.5/HSV1-P53 virus (b). **Confirmation of ΔUL39/Δ34.5/HSV1-P53 construction by PCR (c):** PCR product of the test primers in Δ34.5/HSV-1 indicates an intact band including upstream and downstream sequences of UL39 (lane 3; 671bp), while in ΔUL39/Δ34.5/HSV1-P53, a band (lane 2; 3592 bp) including upstream and downstream sequences of UL39 deleted CDS plus CMV-EGFP-P53-polyA is shown (lane 1, CinnaGen 1 kb DNA ladder).

### Evaluating phenotypic characterization of ΔUL39/Δγ34.5/HSV1-p53 mutant in vitro

The cells infected with ΔUL39/Δγ34.5/HSV-p53 displayed different phenotypes from the parental Δ34.5/HSV-1 (Fig 8). virus and had stronger cell-killing abilities in the tested cell lines that are of various tissue origins and with different p53 status (Vero, BHK-21, A549, MDA-MB-468, Hela, HEK293, HEK293T, Caco-2, and NIH3T3 cell lines). Cultivation of ΔUL39/Δ γ34.5/HSV-p53 mutant in these cell lines was also associated with rounding of cells in early infection without viral cell-to-cell spread and loss of adherence to the monolayer (Fig 9).

**Fig 8.**
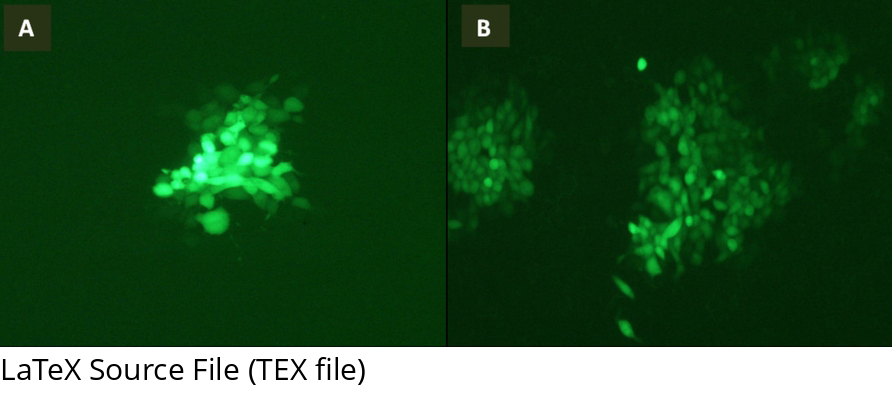
ΔUL39/Δ γ134.5/ HSV1-GFP-infected BHK cells as controls. Fluorescent imaging was done 72 h (A) and 5 days (B) postinfection (Magnification × 100).

**Fig 9.**
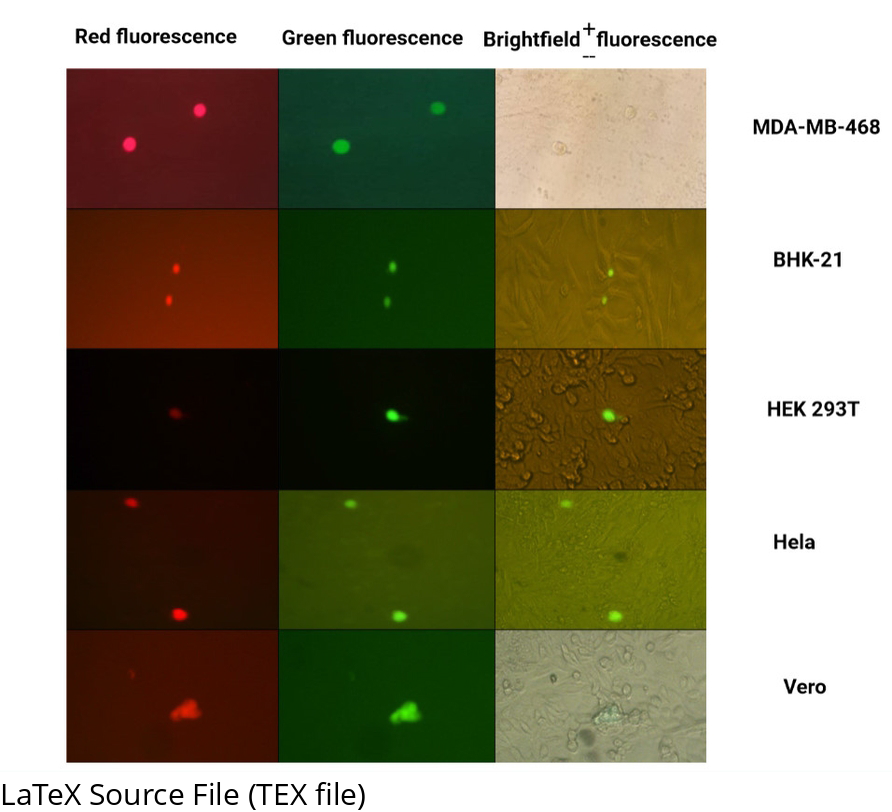
Representative results of cultivation of CAM-adapted HSV-p53 mutant in the cell lines, associating with rounding of cells in early infection without viral cell-to-cell spread and loss of adherence to the monolayer (Scale bar: 200 µm).

## Discussion

In the current study, due to the ability of P53 to trigger apoptosis in target cancer cells, we manipulated the UL39 gene with insertion of the EGFP-p53 expression cassette under the control of the CMV promoter utilizing CRISPR-Cas9 system to improve oncoselectivity and oncotoxicity properties of a neuroattenuated Δγ34.5/HSV-1 mutant.

For isolation of the recombinant ΔUL39/Δγ34.5/HSV-p53, early experiments were examined by the cultivation of the transected/infected viral supernatant in Vero and BHK cell lines that are commonly used in HSV research and then, in cell lines which have shown to use strategies to manage p53 signaling in the favor of their continued survival (MDA-MB-468, Hela, and HEK 293T cell lines). Cultivation of the viral supernatant in these cell lines was also restricted to a single round of replication in cells without viral cell-to-cell spread (Fig 3).

A theoretical explanation may be that the disruption of the UL39 gene (as a viral anti-apoptotic factor) [19] and p53 overexpression during replication of ΔUL39/Δγ34.5/HSV-p53 activate both intrinsic and extrinsic pathways of apoptosis in the virus-infected cancer cells accelerating the premature death, thereby limiting the virus replication to an abortive infection in cells. Hence, we deduced that the defective virus needs to be replicated in a complementary permissive and p53-resistant cellular model [9–11, 43].

In 2012, a few publications reported the ability of pock forming by HSV-1 on the CAM of fertile hens’ eggs in which pocks were white, superficial, and separate that remained typically small [35, 36, 38]. Although the embryonated hen’s egg is largely supplanted by the cultured cells for the isolation and cultivation of viruses, this system is newly used for the in vivo analysis of oncolytic viruses and investigating several functional features of tumor biology such as angiogenesis, cell invasion, and metastasis[44, 36, 35, 33].

However, the highly active metabolism of tissues in a developing embryo in addition to the proliferation of cells in pocks on the virus-infected CAM supplies the essential enzymes for replication of the virus available [38, 39, 44, 45]. Besides, the probable lesser sensitivity of CAM cells of chick embryos to the human p53 effects [15, 46] led us to the hypothesis that the CAM model might be a suitable tissue source for *Δ*UL39/Δ34.5/HSV1-p53 propagation in which the virus can borrow the Ribonucleotide Reductase (RNR) essential enzyme from the host cells and replicate efficiently.

Despite our interesting observations in ΔUL39/Δγ34.5/HSV-p53 multiplication on the CAM, our work highlights the limitations of replication of CAM-adapted HSV-1-p53 in vitro culture. It is suggested that integration of p53 gene into the viral genome makes the cells prone to premature death during virus replication and limits the virus multiplication to a single round of replication resulting in replication-defective mutant viruses production (Fig 10).

**Fig 10.**
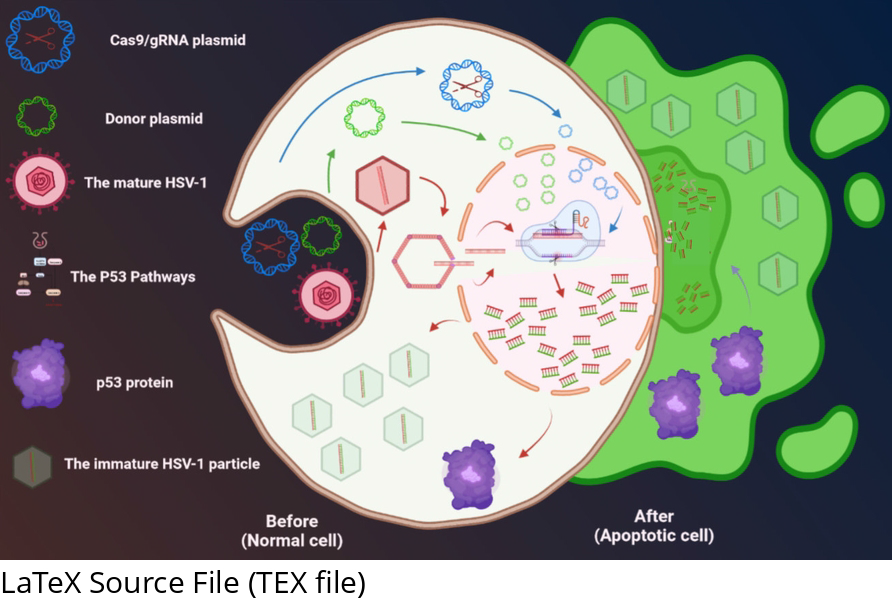
Schematic representation of the fate of cell and virus after during HSV-1 infection after fusion of the virion envelope and host cell membrane, Proteins within the tegument layer of the virus are released into the cytoplasm. The capsid reaches the nucleopores by microtubule transport and releases the viral genome into the nucleus, in where the genome circularizes and sequential transcription of α, β, and γ genes begins. Early in virus replication, p53 gene is overexpressed and balance between the pro- and anti-apoptotic factors is upset and apoptosis is triggered, promoting probably premature death before the virus produces progeny virions. On the other hand, HSV1-P53-induced cell death is often associated with immature virus particles.

Black et al. showed that VSV-encoded p53 inhibited virus replication in non-malignant human pancreatic ductal cells [47]. As well, the finding of other researchers showed the VSV and Adenovirus carrying the p53 gene can simultaneously assist virus replication while enhancing oncolytic potency in cancer cells such as lung carcinoma cells, breast carcinoma cells, cervix carcinoma cells, prostate carcinoma cells, and pancreatic cancer cells [20, 25, 47-50].

Ying et al. reported that the Recombinant Newcastle disease virus expressing P53 (rNDV-P53) has no distinctions in the kinetics and magnitude of replication compared with the vehicle virus (rNDV) in hepatocellular carcinoma model [26].

However, during productive HSV-1 infection, a balance between the pro-apoptotic and anti-apoptotic factors is set up that allows virus propagation. At first, apoptosis is initiated by the immediate early gene expression and later, it is modulated by the early and late viral anti-apoptotic genes which block apoptosis progress [10, 14, 21, 46, 51, 52] .

Although p53 is generally considered an inducer of apoptosis in many viruses-infected cells, previous reports suggested that during HSV-1 infection, p53 plays both positive and negative roles in HSV-1 replication; upregulating ICP27 expression, early during the infection and downregulation of ICP0 at later stages of infection, inhibit apoptosis during the HSV infection. On the other hand, p53’s positive and negative effects in HSV-1-infected cells are organized by multiple mechanisms in a time-dependent and p53 status-dependent manner. However, a threshold of caspase activation must be reached while the balance becomes biased toward cell death through HSV-dependent apoptosis (HDAP). So, it is not surprising that uncontrolled expression of p53 during the virus replication and the increased levels of apoptosis will result in premature host cell death [15–17].

However, the above explanations remains speculative because, to our knowledge, only one HSV1-p53 based study has been reported [53]. Therefore, the results obtained in our study should be interpreted cautiously.

Although the chicken chorioallantoic membrane can be a proper alternative system for isolation of ΔUL39/Δ γ34.5/ HSV-p53, it is necessary to point out the challenges and problems that must be considered for successful use of the CAM model; I) Accurately performing this protocol requires training and practice while the most frequent problem is egg contamination. II) Only the low-titer inoculums of the virus can induce discrete pocks on the CAM, whilst high titer of the virus results in confluent lesions (Fig 4) [38]. So, many eggs may be required for the isolation of recombinant viruses. III) In our research, a drop in viral titer was seen in pock lesions 48 h post-infection, despite cell proliferation progression in the virus-infected CAM, suggesting interferon to be the limiting factor [45]. However, the infection potential of Δγ34.5/HSV-1, as a control virus for CAM was shown to be very different from that seen in cell culture, and the MOIs should be much higher.

In our work, many pocks were required to be collected for infecting a new CAM. On the other hand, the resulting viral titer in each pock is significantly below the required level, while prior studies reported that a single particle of HSV-1 can infect a Vero cell and initiate the replication cycle [54].

To conclude, the CAM can be a promising but challenging model for mass manufacturing of recombinant viruses such as HSV-1-P53 which are not able to replicate in cells.

## Acknowledgements

We hereby thank the staff of Virology Department and Laboratory of Regenerative Medicine and Biomedical Innovations (Pasteur Institute of Iran).

